# The social architecture of an in-depth cellular protein interactome

**DOI:** 10.1101/2021.10.24.465633

**Authors:** André C. Michaelis, Andreas-David Brunner, Maximilian Zwiebel, Florian Meier, Maximilian T. Strauss, Isabell Bludau, Matthias Mann

## Abstract

Nearly all cellular functions are mediated by protein-protein interactions and mapping the interactome provides fundamental insights into the regulation and structure of biological systems. In principle, affinity purification coupled to mass spectrometry (AP-MS) is an ideal and scalable tool, however, it has been difficult to identify low copy number complexes, membrane complexes and those disturbed by protein-tagging. As a result, our current knowledge of the interactome is far from complete, and assessing the reliability of reported interactions is challenging. Here we develop a sensitive, high-throughput, and highly reproducible AP-MS technology combined with a quantitative two-dimensional analysis strategy for comprehensive interactome mapping of *Saccharomyces cerevisiae*. We reduced required cell culture volumes thousand-fold and employed 96-well formats throughout, allowing replicate analysis of the endogenous green fluorescent protein (GFP) tagged library covering the entire expressed yeast proteome. The 4159 pull-downs generated a highly structured network of 3,909 proteins connected by 29,710 interactions. Compared to previous large-scale studies, we double the number of proteins (nodes in the network) and triple the number of reliable interactions (edges), including very low abundant epigenetic complexes, organellar membrane complexes and non-taggable complexes interfered by abundance correlation. This nearly saturated interactome reveals that the vast majority of yeast proteins are highly connected, with an average of 15 interactors, the majority of them unreported so far. Similar to social networks between humans, the average shortest distance is 4.2 interactions. A web portal (www.yeast-interactome.org) enables exploration of our dataset by the network and biological communities and variations of our AP-MS technology can be employed in any organism or dynamic conditions.

---

The large-scale study of cellular interactomes by MS-based proteomics dates back almost 20 years (*1*, *2*), culminating in two studies in which nearly half the expressed yeast proteome was successfully purified with identified interactors (*3*, *4*). These datasets have been mined extensively, leading to a network-based view of the cellular proteome. Given the importance of the interactome for functional understanding and the dramatic improvements in MS-technology during the last decade (*5*, *6*), we set out to generate a substantially complete interactome of all proteins present in an organism in a given state. We made use of an endogenously GFP-tagged yeast library containing the 4159 proteins that were detectable by fluorescence under standard growth conditions (*7*). Miniaturization and standardization of the workflow in combination with an ultra-robust liquid chromatography system with minimal overhead time coupled to a sensitive trapped ion mobility mass spectrometer employing the PASEF scan mode (*8*, *9*), resulted in very high data completeness across pull-downs. This workflow required only 1.5 mL instead of liters of yeast culture, provided a constant throughput of 60 pull-downs per day and allowed using the same conditions for soluble or membrane proteins of vastly different abundances (**Fig. 1A**).

**Figure 1.**
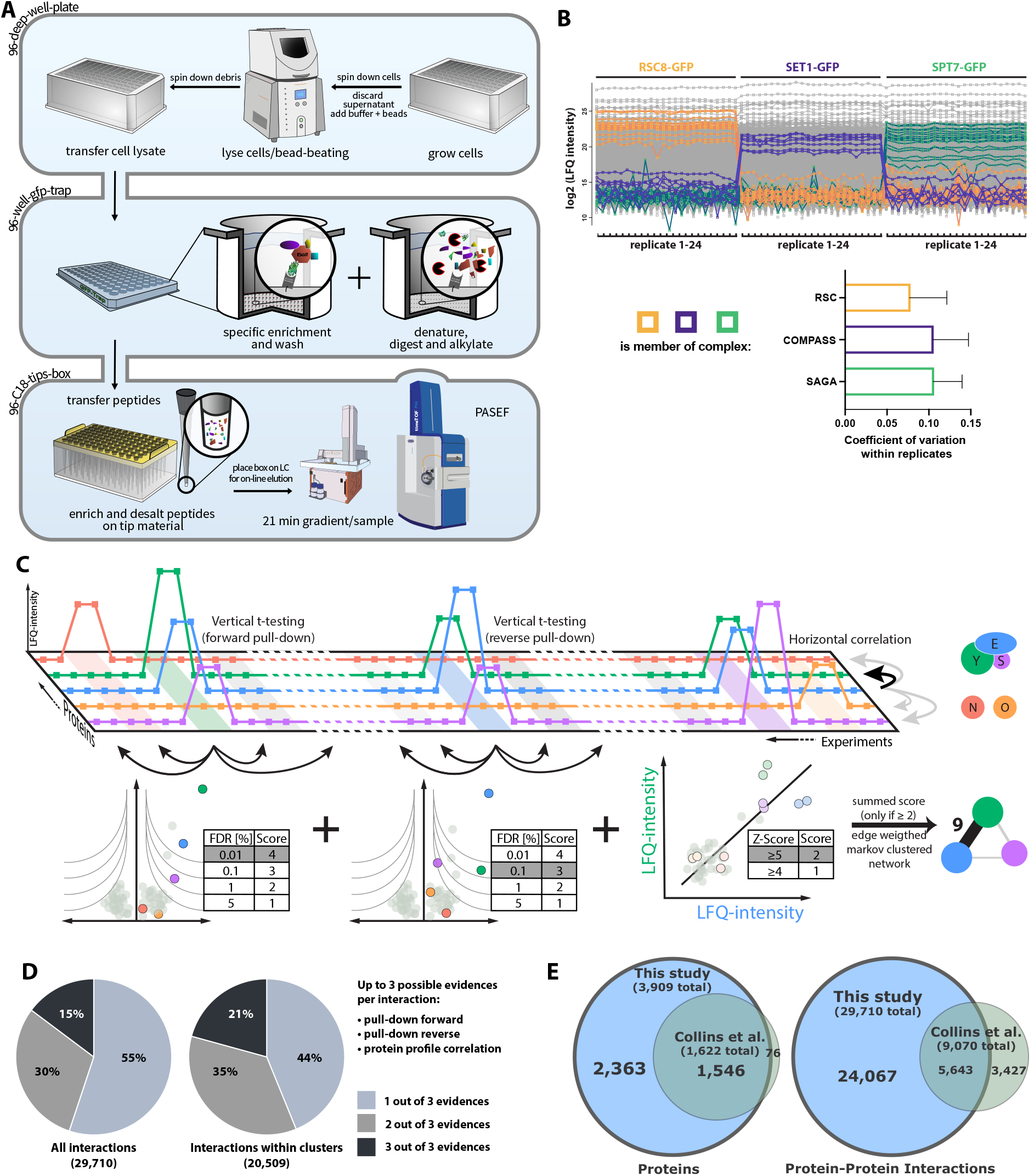
A comprehensive and scalable interactomics technology. **A)** Sample preparation in 96-well format and mass spectrometric measurement: Each strain of the GFP-tagged library is lysed by mechanical disruption and transferred into anti-GFP nanobody coated microtiter plates, where weak interactions are preserved by gentle washing. After enzymatic “in-well” digestion, resulting peptides are transferred on standardized C_18_-StageTips from which they are directly eluted into a standardized 60 samples/day gradient. Data is acquired in the PASEF scan-mode on a trapped ion mobility – Time of Flight mass spectrometer. **B)** Streamlined workflow and reduced transfer steps reduce the risk of manual errors and sample variation: Demonstration of workflow reproducibility and sensitivity on three nuclear complexes in biological replicates. Tagged members of each complex (baits) pull down the known preys in very similar amounts. Lower panel: bar plot of mean coefficient of variation with standard deviations. **C)** Two-dimensional interaction scoring: Columns represent pull-down experiments in replicates (light color). Squares depict intensities of detected proteins across the pull down-experiments. Three levels of evidence support each interaction: t-test of forward pull-down against complement experiments, t-test of reverse pull-down, and protein profile correlation – the correlated abundance profile against all other proteins across all experiments (z-scored, Methods: Protein correlation). **D)** Proportion of interactions backed by multiple layers of evidence. **E)** Overlap of proteins with at least one interactor and interactions detected in this study with the previous state-of-the-art network (*12*).

## Measurement of the yeast interactome

To test the quantitative reproducibility of our workflow, we performed 24 biological replicates of pull-downs of three nuclear complexes, which resulted in complete retrieval of these complexes from a single bait each, with 9% average coefficients of variation (CVs) of enriched complex members (**Fig. 1B**). This compares to a 69% repeatability of assigned interactions in the previous large-scale screens (*10*).

Three layers of evidence help to establish an interaction between two proteins. The first two are statistically significant enrichment of the proteins in the forward and in the reverse pull-downs (where the prey pull-down significantly enriches the bait). Instead of employing only a t-test of bait pull-down against a pull-down of a strain only expressing GFP, we made use of our vast number of diverse GFP-tagged strains, to combine them into a single control group, thereby efficiently removing false positives not specifically binding to the bait (Methods: Enrichment analysis). Using this affinity enrichment (rather than affinity purification) concept (*11*), we quantitatively compared all proteins across more than 8,000 pull-down measurements, making use of the profile similarities of interacting proteins in correlation analysis. This third evidence type turned out to be very informative due to the large quantitative accuracy combined with close to a complete set of “virtual controls” (Methods: Protein correlation, **Fig. 1C**).

We combined all three layers of each interaction into a single interaction score and retained those with a minimum score of 2, corresponding to (a) a single pull-down at 1% FDR or (b) a correlation z-score of at least five or (c) forward and reverse pull-downs at 5% FDR each, or (d) one at 5% FDR combined with a correlation z-score greater than four. To retrieve clusters and complexes from our interactome data, we used Markov clustering with the above-derived score as the edge weights, without any training or a priori knowledge (Methods: Network generation, **Fig. 1C**).

The replicate GFP pull-down measurement in the 4,147 yeast strains resulted in the enrichment of 82% of the baits (**Suppl. Fig. 1**). Our MS-data provided statistically significant evidence for a total of nearly 30,000 physical interactions, corresponding to an average of 15.2 interactions per protein. Most were supported by forward pull-down (38%), followed by forward pull-down and significant prey correlation (29%), whereas nearly all interactions with both forward and reverse evidence also had significant correlations (> 99%) **(Suppl. Fig. 2)**.

Due to the limited overlap of the interactions reported by two previous large-scale studies (13% shared interactions), *Collins et al*. merged and reanalyzed these datasets to create a consensus network with 1,622 nodes (*12*). Our data encompasses 95% of these, but places nearly the entire expressed yeast proteome in a network (3,909 nodes). Our dataset of 30,000 significant protein-protein interactions confirms 62% of the much smaller *Collins et al*. dataset (**Fig. 1E**). Based on a comparison with the BioGRID database (*13*), over two-thirds of the interactions reported here are novel.

## Organization of protein-protein interactions in clusters

Markov clustering analysis - with our interaction scores as edge weights, condensed the network into 623 clusters, with about 20,000 interactions within them, most supported by at least two statistically significant levels of evidence (**Fig. 1D**). When we inspected known protein complexes from different cellular compartments, especially membrane complexes, we found them to recapitulate the literature to a large degree. Furthermore, we here retrieved 3628 interactions between membrane annotated proteins, compared to 853 in a dedicated membrane proteome (*14*). This is shown exemplarily for the full retrieval of the endosomal retromer complex, the conserved oligomeric Golgi complex, and the plasma membrane exocyst complex (**Fig. 2A**). At the same time, our unbiased and high coverage analysis identified novel subunits with tight association to known complexes. For instance, three subunits of the essential endoplasmic reticulum (ER) membrane oligosaccharyl transferase (OST) complex - an integral component of the translocon - associated with α-1,2-mannosidase (Mns1; human homolog: MAN1B1), an enzyme that catalyzes the ER glycoprotein trimming reaction which is required for ER-associated protein degradation (ERAD). This indicates that the enzymatic activity of N-linked oligosaccharide chain addition is physically connected to the removal of a terminal sugar, at least in one isoform of the OST complex. The slow enzymatic activity of Mns1 acts as a timer (*15*, *16*) and we speculate that it co-translationally primes stalled or erroneous proteins directly at its site of translocation for ERAD degradation. We also discovered a novel complex defined by three unreported interactions (all with the maximum interaction score of 10) between Tcd1, Tcd2 - mitochondrial proteins that are involved in tRNA base modification - and YGR012W, a protein of unknown function. A homolog of Tcd1 and Tcd2 in *E. coli* termed TcdA functions in a complex of three in the cyclization of an essential tRNA modification found in all three domains of life (*17*).

**Figure 2.**
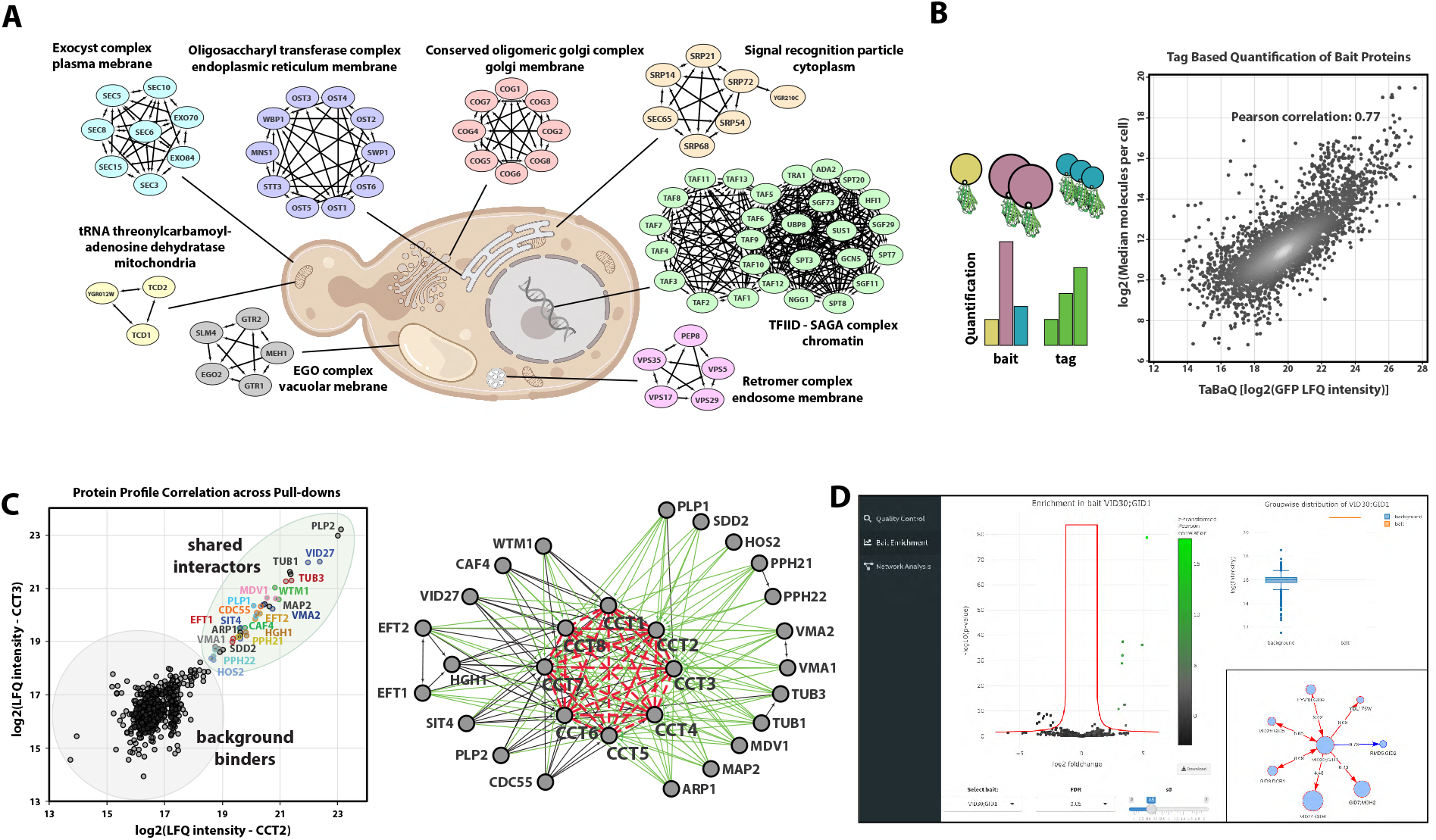
High-quality dataset for the exploration of the interactome. **A)** Clusters derived from our interactome for a range of challenging complexes such as chromatin-associated, soluble and membrane-bound complex of various organelles. In each case, all known subunits were retrieved. **B)** Tag-based quantification allows retrieving abundance information for the baits in a generic manner (left panel). Correlation of tag peptide-based signals with a literature compilation of yeast protein abundances (*18*) (right panel). **C)** For the non-taggable chaperonin containing t-complex (CCT), profile correlation analysis nevertheless reveals its subunits and interactors. Interactions based on correlation only are shown in red (dashed) and unreported interactions with CCT in green. **D)** Web application that allows exploration of interaction data for interactions of interest. For all proteins, pull-downs are depicted as volcano plots together with a violin plot that shows the MS intensity of user-selected outliers. Subnetwork from pull-downs of the selected bait and reverse pull-downs or significant interactors.

Many biological complexes share members and these can be difficult to disentangle by clustering algorithms. We speculated that our highly quantitative data could nevertheless resolve these cases. Applying a network layout algorithm (Methods: Network generation) to members of the transcription factor TFIID and the SAGA complex, separately reconstructed these complexes, while correctly assigning shared members (**Fig. 2A**). At the global scale, we found that about two-thirds of all interactions connected members within clusters, whereas the remainder connected clusters to each other. For example, the cytoplasmatic signal recognition particle (SRP) is connected to another cluster containing the SRP-receptor (SRP101/102). The largest connected clusters were the small and large subunits of the ribosome, with 362 inter-complex connections.

Leveraging the common, endogenous GFP-tag on more than 3379 detected baits, we next investigated if the MS-signal of the GFP peptides could be used to quantify each bait. Indeed, these intensities correlated well (r = 0.77), with a recent compilation of yeast protein abundances (*18*) (**Fig. 2B**). This validates our interaction workflow and allows tag-based estimation of the relative abundances of proteins in a cluster, which is useful to determine their functional role (*19*).

For some proteins, for example the members of the chaperonin containing t-complex (CCT), tagging is not possible because it interferes with protein stability or function (*20*). Based on highly significant correlations between profiles of the subunits, CCT was nevertheless fully recovered (**Fig. 2C**). Besides the eight conserved, ring-forming members, we also detected a distinct set of 21 interacting proteins, about half of which had not been reported yet. Two of these were catalytic subunits of protein phosphatase 2A, suggesting regulatory functions, and others, such as tubulin and actin-related proteins (Tub1, Tub3, Arp1) major known folding substrates. CCT may have a restricted or broad set of folding substrates (*21*), and our results quantitatively support the former possibility.

The above examples only scratch the surface of the interesting biological leads contained in the data. To allow ready exploration of interactions of interest, we created a web portal (www.yeast-interactome.org), which supplies statistical evidence for protein-protein associations, and summarizes the resulting clusters (**Fig. 2D**).

## Network architecture of the cellular interactome

The availability of data for large networks in systems ranging from power-grids, genetic networks to human social networks, has enabled the study of their underlying architecture, commonalities and differences (*22*). This topic also has a long history in protein interaction networks. However, these analyses have been limited by the incompleteness of the data, especially in multicellular species (*23*). With an in-depth protein-protein interaction map in hand, we compared its characteristics to networks in different domains. Yeast proteins are highly connected with an average of 15 and a median of 6 interactions per protein, significantly more than the human BioPlex interactome (average interactions: 8) (*24*) (**Fig. 3A**). Influential nodes – those with the highest number of normalized interactors (or degree centrality) – were more common than in the GitHub package dependency network, but less common than in a similarly-sized Facebook subnetwork (**Suppl. Fig. 4**). This high connectivity is reflected in a mean shortest path between yeast proteins of only 4.2, ranging from highly connected proteins with only three steps to less connected ones with an average of more than 7. (**Fig. 3B**). This is very similar to the 4.7 path-length for world-scale Facebook relationships (*25*).

**Figure 3.**
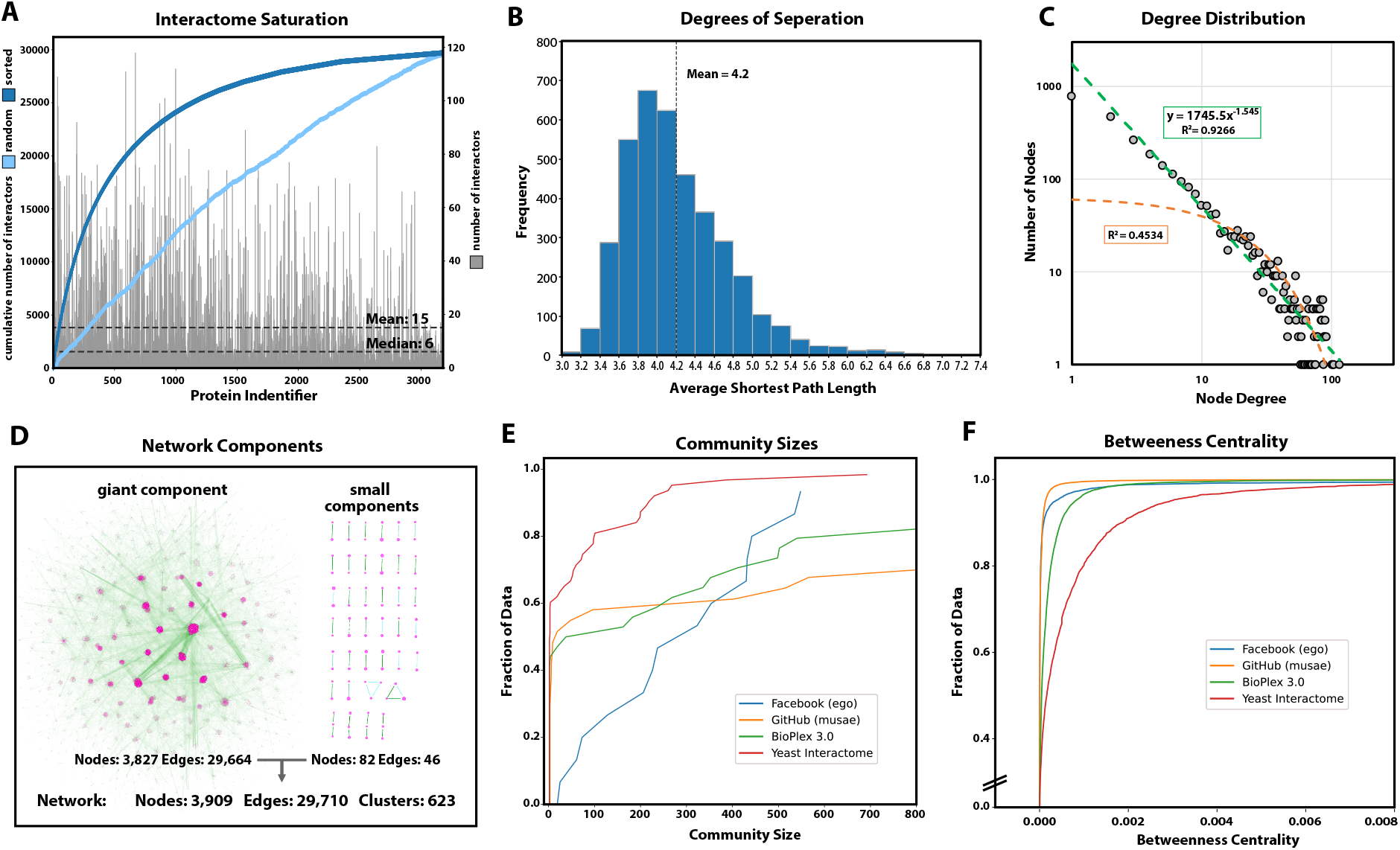
Properties of the protein interaction network. **A)** Distribution of the number of interactors (grey). Sorted cumulative number of interactions reaches saturation at 30,000 interaction (blue) **B)** The distribution of average shortest path length between all possible pairs of nodes within the giant component shows a mean of 4.2 steps corresponding to 3.2 intermediaries (“degrees of separation”) **C)** Power-law fit (green; equals a linear fit on a log-log scale) of the frequency of proteins with a given number of interactions highlights the scale-free properties of the network. Exponential fit depicted in orange **D)** Nearly all nodes of the network are connected with each other in the giant component. **E)** Cumulative distribution function of the community sizes (Louvain algorithm) detects more smaller communities for *S. cerevisiae*. **F)** Cumulative distribution function of betweenness centrality: The *S. cerevisiae* interactome has more nodes with a high betweenness-centrality than the comparison data sets.

One of the key features for most real-life networks with complex topology in contrast to random networks is the scale-free power-law distribution of interactors (*26*, *27*). Scale-free network properties are thought to arise by preferential attachment over evolutionary time to already well-connected nodes and can be identified by a linear relation of the node degree or number of interactors with its frequency (number nodes with that degree) plotted in log-log space. While this has been hard to prove for biological networks, they rather appear to be exponential or have a truncated power-law degree distribution (*28*), our yeast interactome clearly displays scale-free properties (**Fig. 3C**). In accordance with previous protein-protein interaction networks (*3*, *29*), the exponent was below two, at the lower end of the two to four range of other scale-free networks.

The high connectivity of most proteins organizes almost all of them (3,827) into a single giant connected component, accompanied by 38 small components (82 proteins) (**Fig. 3D**). A total of 478 proteins were outside of the network because MS-analysis of their pull-downs only identified the bait itself. There was an significant enrichment for 87% of these baits (FDR<0.01%), indicating that there were no identifiable interactors under our standard conditions despite a successful pull-down (**Suppl. Fig. 3**, see volcano plots accessible via web-application).

We next investigated the large-scale organization of the yeast interactome using the Louvain community detection algorithm (Methods: Network comparisons). This revealed that yeast is organized in smaller communities than GitHub, ego-Facebook and also Bioplex (**Fig. 3E**). Important “bottleneck” proteins that are part of many shortest paths have a high “betweenness-centrality”. The yeast interactome has comparably more of those central nodes and bioinformatic enrichment analysis highlighted proteins involved in “RNA polymerase II”, “mitochondrial nucleoid”, “gluconeogenesis” and “misfolded protein binding” (**Fig. 3F; Suppl. Table 1**).

Altogether, based on the total of 4,387 identified yeast proteins, only 10.9% had no discernable interaction partner, whereas 74.2% had at least two. Given that some of our baits will have context dependent interactions not captured here, our estimates are conservative and we conclude that almost all yeast proteins are “social”.

## Global organization in clusters highlights novel interactions

Intensive research over the last decades has made *S. cerevisiae* arguably the best understood single-cell eukaryotic organism, leading to the discovery of crucial conserved cellular functions, such as metabolic pathways, mechanisms of DNA replication and transcription, protein quality control and modifications that were later confirmed in human and other organisms. Nevertheless, our interactome still contained uncharacterized proteins or interactions not reported in the BioGRID database and thus providing novel biological insights (extended selection **Suppl. Fig. 6**). Furthermore, BioGRID has accumulated binding events from very disparate experiments without a common confidence score (133,900 physical interactions from about 10,000 publications). We reasoned that our homogeneous, high-quality data set would help biologists to highlight true positive interactors with biological relevance, several of whom we discuss below.

A total of eleven evidences connect the uncharacterized protein YDL176W with the conserved glucose-induced-degradation (GID) complex, only a few of which had been indicated by previous pull-down or genetic interaction data (*3*, *30*) (**Fig. 4B**). These types of high-confidence associations assist in prioritizing interactions and form the basis for a detailed mechanism and structure discovery of a novel GID modulator. Similarly, our data ties the uncharacterized protein YJR011C to the conserved transcription and translation regulatory CCR4-Not complex (*31*, *32*) via high-significant interactions to a majority of its subunits (**Fig. 4G**). Finally, YHR131C is linked to three and YLR407W to the fourth subunit of the kinase CK2 (**Fig. 4N**). We discovered an interaction of Cue4 – a protein of unknown function containing a ubiquitin-binding domain – with the ER membrane complex EMC, potential membrane protein chaperone (**Fig. 4L**). As Cue4 is a paralogue of Cue1 (coupling of ubiquitin conjugation to ER degradation), a component of ERAD (*33*), this physical link and the known aggravating genetic interactions of *Δcue1* with EMC knock-outs (*34*) suggests an ERAD related quality control mechanism for EMC.

**Figure 4.**
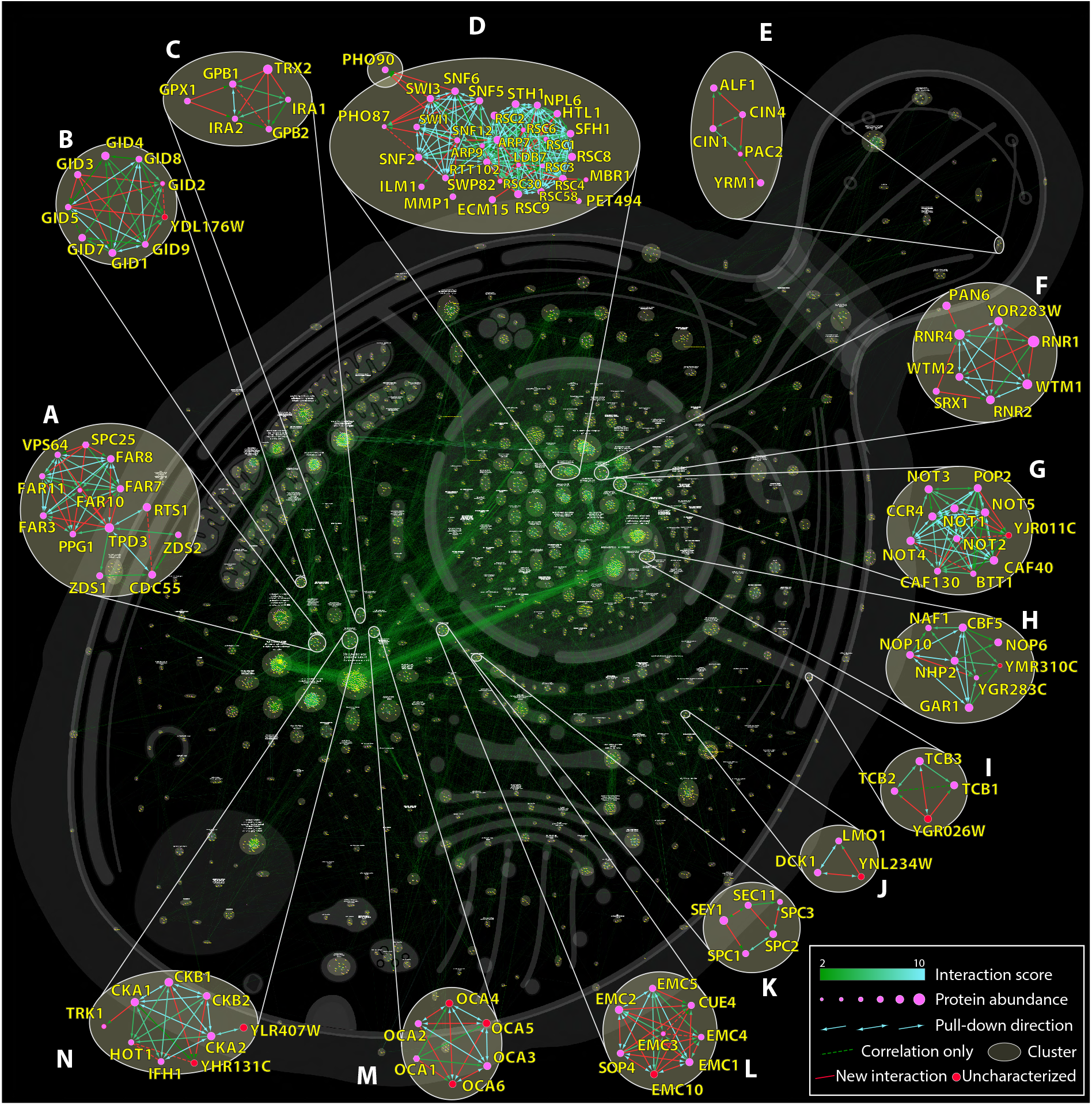
Network of an in-depth interactome highlighting novel interactions. Cellular interaction map of all significant interactions. Clusters are highlighted by circles and cellular localization is indicated by most frequent GO term within a cluster. Enlargements show examples of either novel interactions (based on BioGRID) or those that have not been described further as potential high significant interactor and interactions involving uncharacterized proteins. A full browsable and interactive version of this network can be found at our web application (www.yeast-interactome.org).

The transcriptional regulator SWI/SNF unexpectedly interacts with the phosphate transporters Pho87 and Pho90 (**Fig. 4D**). Out of four plasma membrane phosphate transporters only Pho87 and Pho90 comprise a cytoplasmatic accessible SPX domain. While an SPX dependent phosphate sensing mechanism has been discovered in plants (*35*), it remains elusive in *S. cerevisiae*. In *Arabidopsis* inositol pyrophosphate InsP_8_ concentration increases under phosphate rich conditions and promotes the interaction between SPX domains and a four-stranded coiled-coil motif of phosphate starvation response transcription factors (*36*). Strikingly the recently solved structure of SWI/SNF reveals such a coiled-coil four-helix-bundle at its spine region (*37*) providing a potential SPX interaction site. This raises the possibility of a novel cytoplasmatic sensing and retention mechanisms of this key transcriptional regulator which is known to be necessary for a phosphate starvation response (*38*, *39*). Interestingly, not only the SWI/SNF complex but also an SPX domain-containing phosphate transporter named XPR1 - which has recently been shown to be controlled by InsP_8_ (*40*) - is present in humans.

Illustrating translational relevance, we expand the known interaction of the GTPase-activating protein Ira1/Ira2 (NF1/neurofibromin in humans) and Gpb1/Gpb2 (ETEA in humans) (*41*) by Trx2 a thioredoxin isoenzyme (human homolog: TXN) and Gpx1 (human homologs: GPX3-6), an antioxidant enzyme whose glutathione peroxidase activity is neuroprotective in models of Huntington’s disease (*42*) (**Fig. 4C**).

Additionally, we find a new physical interaction between the two uncharacterized proteins YPR063C and YNR021W (**Suppl. Fig. 6**) whose dimerization and structure has just been predicted in a deep-learning approach (*43*).

Apart from known and novel protein complexes, the yeast interactome depicted in **Fig. 4**, clearly shows evidence of high order connections. These often map to different compartments of the cell, such as the prominent connections between ribosomes in the cytoplasm and the nucleolus, its site of maturation or connect large and small ribosomal subunits that despite its “stickiness” are organized in individual clusters.

## Outlook

Here we have developed and applied a novel and highly scalable interactome technology, enabling replicate measurement of the yeast network in a fraction of the measurement time and starting materials needed previously. Our screen reached near saturation and contained nearly all complexes expected under our experimental conditions (**Fig. 3A, Fig. 4**). Given its streamlined nature, our workflow can now readily be used in other endogenously tagged model organisms (*44*) or to study remodeling of the interactome in the presence of dynamic biological processes or perturbations. Similarly, we envision its use with other interaction technologies like BioID or APEX using tagged libraries that nowadays can be easily generated using the SWAp-Tag platform (*45*). The comprehensive yeast interactome data can further be used as prior knowledge for hypothesis-driven analysis of protein complexes, for example for native protein complex co-fractionation coupled to MS (*46*, *47*). Additionally, we imagine that such interactome data could also be combined with MS-crosslinking studies and recent advances in computational prediction of protein structures from their sequences (*48*, *49*) to yield complete structural models in many cases.

## Acknowledgments

We thank the former members of the Stefan Jentsch lab for their support in yeast cell culture related issues, especially Jochen Rech for providing and handling the library. We greatly appreciate discussions with Marco Y. Hein and Fabian Hosp who were sharing their experience with large-scale pull-down screens and Brenda Schulman for project guidance and discussions. We thank all members of the Department of Proteomics and Signal Transduction for help and discussions and in particular Igor Paron and Christian Deiml for mass spectrometry assistance, Mario Oroshi for technological assistance and Medini Steger for manuscript proofreading. We thank our core facility for their great services and Bruker, Chromotek and Evosep for their continuous technical support.

## Experimental Methods

### Cell growth

To achieve samples with similar cell numbers, pre-cultures of the *S. cerevisiae* GFP-tagged library were grown in YPD media (1% yeast extract, 2% bacto™ peptone, 2% glucose) for two days in 2 mL, u-bottom shaped 96-deep-well plates. This allowed cell concentration convergence of different strains during the slow growing post-exponential phase. Cells were resuspended and 50 μl of each pre-culture was used to inoculate 1.5 mL of fresh YPD media (corresponding to an optical density of 0.5 at 600 nm) in 96-deep-well plates (LoBind®, 2 mL, cat no. 0030504305, Eppendorf AG, Hamburg, Germany). Plates were covered with an air permeable membrane and incubated while shaking at 300 rpm and 30 °C for 6 hours. This allowed the progression through the lag phase and three cell cycles followed by harvesting under standard growth conditions. Cells were pelleted in the 96-deep-well plates by centrifugation at 3500 rpm (= 2451 g) for 5 min. The supernatant was discarded by fast decanting and quick dabbing on paper towels. Plates with pellets were sealed with plastic covers and stored at −80 °C until cell lysis.

### Cell lysis

Dee-well plates with cell pellets were thawed on ice for 5 min. 100 μl of glass beads (0.5 mm, acid-washed, cat no. G8772, Merck KGaA, Darmstadt, Germany) were added to each well using a 96-well bead dispenser (LabTIE International, Veenendaal, Netherlands). After 5 min 250 μl of 4 °C cold lysis buffer (50 mM Tris HCl pH 7.5, 150 mM NaCl, 5% glycerol, 0.05% IGEPAL CA-630, protease inhibitor EDTA-free (cOmplete™, 1 tablet per 50 mL, cat no. 11873580001, Merck KGaA, Darmstadt, Germany), 1 mM MgCl_2_, 0.75 U/μL in-house *Serratia marcescens* endonuclease/SmDNase) were added. Plates were sealed using a heat sealer (S200, cat no. 5392000005, Eppendorf AG, Hamburg, Germany), the low profile plate adapter (cat no. 5392070020, Eppendorf AG, Hamburg, Germany) and transparent heat sealing films (cat no. 0030127838, Eppendorf AG, Hamburg, Germany) for 2 sec at 180 °C and immediately put back on ice. Cell lysis was performed within the 96-deep-well plates at 4 °C via bead-beating (2010 Geno/Grinder®, SPEX SamplePrep, Metuchen, NJ) for 4 cycles of 1.5 min each at 1750 rpm. Plates were cooled in ice water and covered with ice for 7 min in-between cycles and for 10 min after the last cycle. 4 plates were processed in parallel during bead-beating and top and bottom positions were switched at each cycle. Cell debris was spun down at max speed (4300 rpm = 4347 g) for 10 min at 4 °C. Plates were carefully put back on ice and immediately used for the pull-down protocol (Fig. 1A).

### Interactor enrichment

Pull-downs and all sample handling steps were performed at 4 °C. Anti-GFP nanobody coated 96-well microtiter plates were custom made and optimized for this protocol allowing efficient and high reproducible “in-well” digestion, and mass spectrometry compatibility (plates are now commercially available as: GFP-Trap® Multiwell Plate, cat no. gtp-96, Chromotek GmbH, Martinsried, Germany). Plates were prepared with 200 μL wash buffer 1 (50 mM Tris HCl pH 7.5, 150 mM NaCl, 5% glycerol, 0.05% IGEPAL CA-630) per well on a shaker for 1 min at 800 rpm followed by removal of the buffer. The cell lysates were carefully transferred from the 96-deep-well plates by slow uptake of 175 μL supernatant without dislodging glass beads nor the cell debris pellet to the GFP-Trap plate. The GFP-Trap plate was incubated for 1 h at 800 rpm on a small stroke (3 mm) shaker (TiMix 5 control, Edmund Bühler GmbH, Tübingen, Germany) to enrich for GFP-tagged proteins and their interactors. Cell lysates were discarded and plate wells were washed twice with 200 μL wash buffer 1 and twice with wash buffer 2 (50 mM Tris HCl pH 7.5, 150 mM NaCl, 5% glycerol). To allow stable binding of unspecific background proteins – an important factor for label-free quantification – wash buffer was added slowly, and plates were not shaken during wash steps. Emptied, protein-enriched plates were covered and stored at −80 °C until mass spectrometry sample preparation (Fig. 1A).

### Sample preparation for mass spectrometry

Protein-enriched GFP-Trap plates were brought to room temperature and 50 μL of digestion mix 1 (4.5 M urea, 1.5 M thiourea, 10 mM Tris HCl pH 8.5, 3 mM dithiothreitol, 2 ng/μL LysC) were added per well. Plates were incubated at 30 °C and 1000 rpm on a small stroke (3 mm) shaker. After 3 h, 100 μL of digestion mix 2 (10 mM Tris HCl pH 8.5, 7.5 mM chloroacetamide, 2 ng/μL LysC) were added and microtiter plates and lids were sealed with parafilm®. The plates were incubated overnight at 30 °C/800 rpm. The reaction was stopped and the sample was acidified with 15 μL of 10% TFA per well. Plates with peptides were stored at −80 °C till sample loading on EvoTips (Evosep, Odense, Denmark) (Fig. 1A).

### Loading of peptide samples on Evotips

Evotips (Evosep, Odense, Denmark) were activated for 5 min in a 1-propanol Evotips-box reservoir at room temperature (RT), followed by a wash step with 50 μl buffer B (acetonitrile (ACN) with 0.1 % formic acid (FA)) and centrifugation at 500 g for 1 min at RT. The flow-through was discarded and Evotips were placed back into 1-Propanol. Evotips were conditioned with 50 μL of buffer A (ddH_2_O with 0.1 % FA) and centrifugation at 500 g for 1.5 min at RT and were placed in a container with buffer A. 40 μL of thawed peptide sample were loaded and Evotips were centrifuged at 500 g for 1.5 min at RT and placed back in a container with buffer A. 200 μL of buffer A were added and partially washed through the Evotips by centrifugation at 500 g for 50 s. Evotips boxes with buffer A at the container bottom were placed on the Evosep One liquid chromatography (LC) platform (Evosep, Odense, Denmark) for LC-MS analysis. Pull-downs were acquired in technical duplicates and the injection order was reversed after the first measurement (Fig. 1A).

### Liquid-chromatography

For separating peptides by hydrophobicity and eluting them into the mass spectrometer, we used the EvoSep One LC system and analyzed the yeast interactome pull-down proteomes with the standardized 21 min (60 samples per day) gradient. We employed a 15 cm × 150 μm inner diameter column with 1.9 μm C18 beads (PepSep, Marslev, Denmark) coupled to a 20 μm ID electrospray emitter (Bruker Daltonik GmbH, Bremen, Germany). The column was replaced between replicate measurements. Mobile phases A and B were 0.1 % FA in water and 0.1 % FA in ACN, respectively. The EvoSep system was coupled online to a trapped ion mobility spectrometry quadrupole time-of-flight mass spectrometer (*50*) (timsTOF Pro, Bruker Daltonik GmbH, Bremen, Germany) via a nano-electrospray ion source (Captive spray, Bruker Daltonik GmbH, Bremen, Germany). A 24-fraction library of wild-type *S. cerevisiae* was generated using the high-pH reversed-phase “spider-fractionator” (*51*) and data were acquired using the same sample set-up.

### Mass spectrometry

Mass spectrometric analysis was performed in a data-dependent (dda) PASEF mode. For ddaPASEF, 1 MS1 survey TIMS-MS and 4 PASEF MS/MS scans were acquired per acquisition cycle. The cycle overlap for precursor scheduling was set to 2. Ion accumulation and ramp time in the dual TIMS analyzer was set to 50 ms each and we analyzed the ion mobility range from 1/K_0_ = 1.3 Vs cm^−2^ to 0.8 Vs cm^−2^. Precursor ions for MS/MS analysis were isolated with a 2 Th window for m/z < 700 and 3 Th for m/z >700 in a total m/z range of 100-1,700 by synchronizing quadrupole switching events with the precursor elution profile from the TIMS device. The collision energy was lowered linearly as a function of increasing mobility starting from 59 eV at 1/K_0_ = 1.6 VS cm^−2^ to 20 eV at 1/K_0_ = 0.6 Vs cm^−2^. Singly charged precursor ions were excluded with a polygon filter (otof control, Bruker Daltonik GmbH, Bremen, Germany). Precursors for MS/MS were picked at an intensity threshold of 2,000 arbitrary units (a.u.) and re-sequenced until reaching a “target value” of 24,000 a.u. considering a dynamic exclusion of 40 s elution. The capillary voltage was set to 1,750 V and dry gas temperature to 180 °C.

### Raw data processing

MS raw files were processed using MaxQuant (v1.6.17.0) (*52*, *53*), which extracts features from four-dimensional isotope patterns and associated MS/MS spectra, on a computing cluster (SUSE Linux Enterprise Server 15 SP2) utilizing UltraQuant (github.com/kentsisresearchgroup/UltraQuant). To allow processing in an acceptable time frame, RAW files were handled in 5 parallel batches of approximately 1700 files each containing plates equally distributed across the measurement period. Files were searched against the *S. cerevisiae* Uniprot databases (UP000002311_559292; canonical and isoform, reviewed-sp and unreviewed-tr from 02/2020). For high significance identification the false-discovery rates were reduced and controlled at 0.1% both on peptide spectral match (PSM) and protein levels. Peptides with a minimum length of seven amino acids were considered for the search including N-terminal acetylation and methionine oxidation as variable modifications and cysteine carbamidomethylation as fixed modification, while limiting the maximum peptide mass to 4,800 Da. Enzyme specificity was set to LysC cleaving C-terminal to lysine. A maximum of two missed cleavages were allowed. The parameter “type” was set to “TIMS-DDA” with “TIMS half width” at 4. The instrument was set to “Bruker TIMS” and main search peptide tolerance reduced to 8 ppm, the max. charge set to 5 and min. peak length to 3. Peptide identifications by MS/MS were transferred by matching four-dimensional isotope patterns between the runs (4D-MBR) using a narrow elution match time window of 12 s and a reduced ion mobility window of 0.01 1/K_0_. Protein quantification was performed by label-free quantification using a minimum ratio count of 2. The 24-fraction library was added as an additional parameter group with the same group-specific settings, but LFQ disabled and “separate LFQ in parameter groups” under global parameters enabled. The writing of additional tables was disabled for performance reasons.

### Raw data availability

All mass spectrometry raw data and MaxQuant output tables have been deposited to the ProteomeXchange Consortium (*54*) via the PRIDEpartner repository with the dataset identifier available upon publication.

### Data processing and normalization

Twelve outdated samples of the GFP library were eliminated. These included wrongly annotated ORFs that were merged with others: YAR044W, YPR090W, YDR474C, YFR024C, YJL021C, YJL017W, YGL046W, YFL006W, YGR272C, YBR100W, YJL018W, YJL012C-A. After the removal of potential contaminants, reverse and “only identified by site” hits, MaxQuant proteinGroups.txt output files from the 5 batches were merged using the majority protein IDs column. Values were filtered for two valid values within at least one replicate group. To adjust for potential differences between the 5 MaxQuant batches caused by the parallel applied label free normalization algorithm and for potential handling batch effects between 96-well plates, values were median normalized if there were more than 5% of valid values in each of the corresponding groups.

### Missing value imputation

Missing values were imputed in a two-tiered approach. For proteins with measured values in more than 5% of all samples (or minimally 400 samples), a protein-specific missing value imputation approach was used. Here, a random value was sampled from a normal distribution with following properties: mean = median of all measured intensity values for the given protein, standard deviation = standard deviation of all measured intensity values for the given protein. Lower and upper bounds for the normal distribution were set to three standard deviations from the mean and minimally to zero. The function “rtruncnorm” from the R library “truncnorm” was employed. For proteins with less than 5% valid values (or in less than 400 samples), global metrics were employed for missing value imputation. Here, missing values were sampled from a normal distribution with the following parameters: mean = mean of all quantified values across all proteins and samples minus 1.8 times the standard deviation, standard deviation = the standard deviation of all quantified values across all proteins and samples multiplied by 0.3.

### Protein correlation

Due to the large sample number that would negatively influence correlation, we chose a subsampling approach: For each protein pair across the sample profile, the top 2% of samples with the highest intensities for both proteins were selected (resulting in 2-4% depending on their overlap) and complemented by twice the number of randomly selected samples as background. The selected subset of samples was used to calculate the Pearson correlation coefficients of the protein pair (Fig. 1C). The effect of weighted correlation can be visualized by enabling “subsample values” under protein correlation in our web application (yeast-interactome.org). Since the distributions of correlation coefficients varies between proteins and in order to define a universal cut-off for significant correlations, correlation coefficients were normalized via row wise z-scoring. A z-scored Pearson correlation coefficient above 4 and 5 therefore corresponds to a chance probability of below 3.2*10^−5^ and 2.9*10^−7^, respectively.

### Enrichment analysis

A two-tailed Welch’s t-test was performed on each replicate-grouped pull-down sample using all corresponding complement samples as a combined control (*11*). Within the combined control group, samples with the highest bait correlation (top 5%) were excluded in order to provide a bait-unrelated control. FDR cutoff-lines were calculated using an analytical approach using an S0-parameter of 0.5 (*55*).

### Network generation

Interactions for the first two layers of evidence (forward and reverse pull-down) were defined between bait proteins and significantly enriched prey proteins from the t-tests. They were scored based on their FDR of 5%, 1%, 0.1% and 0.01% at 1, 2, 3 and 4, respectively (“score_FDR”). For the third layer of evidence, an interaction for z-scored Pearson correlation coefficients above 4 and 5 was scored at 1 and 2, respectively (“score_cor”). All three layers of evidence were combined into a single interaction score ranging from 1-10 (“score_FDR+cor”), thereby weighting interactions based on their experimental significance (Fig. 1C). Networks were created and exported into Cytoscape (*56*) for further analysis and visualization strategies. The network was filtered for interactions with a combined score equal to or above 2, thereby excluding interactions based only on a single t-test with an FDR of above 1% or a z-scored Pearson correlation coefficient of below 5. The Markov clustering algorithm was applied using the interaction score as edge weight and a granularity parameter of 2.5 while retaining inter-cluster edges. The “CompoundSpringEmbedder” (CoSE) layout algorithm was applied to single clusters. The network including edges (interactions) and nodes (proteins), annotations, and layouts can be downloaded as Cytoscape session at (www.yeast-interactome.org) or at the NDEX network database (*57*) via the UID available upon publication.

### Organelle based mapping of clusters

Within the Cytoscape group preferences the attribute aggregation was enabled and “visualization for group” were set to “none”. The WordCloud “minimum word occurrence” and the “max. words per label” was set to 1, and normalization to 0. To generate outcome with location specific words only, the excluded words list was extended by following terms: apparatus, matrix, membrane, intermembrane, chromosome, ii, protein, anchor, coated, cytoplasmic, iv, lipid, pass, peripheral, secreted, pit, side, single, centromere, type, endomembrane, tip, reticulum, body, localizes, kinetochore, gpi, note, neck, prospore, granule, replication. The “AutoAnnotate” plugin (*58*) was used to generate localization-based name for each markov cluster utilizing WordCloud (*59*) (most abundant word within “Subcellular localization [CC]”). Collapsed localization (collapse singleton clusters enabled) based labeled groups were organized using the “Boundary Layout” using self-defined areas. Node repulsion was increased to 1,000,000. For cluster annotation the standard complex name from EMBL Complexportal was used. For each cluster the two most frequent names were used, (minimum word occurrence: 2). The image of the background cell in Figure 4, the Cytoscape session and the web application is an adopted version from SwissBioPics by the Swiss-Prot group of the SIB Swiss Institute of Bioinformatics. Cell image in Figure 2A was created with BioRender.com.

### Network comparisons

Network comparison analysis was performed in Python 3.8.1. Tabular data was loaded via the pandas package (1.3.1) and converted to a network via NetworkX (2.6.2). To calculate “Betweeness” and “Degree Centrality”, the respective NetworkX functions were used. To perform community analysis, a Python implementation of the Louvain algorithm was used (https://github.com/taynaud/python-louvain, version 0.15). Cumulative distribution functions were plotted using the matplotlib-library (3.4.2) and NumPy (1.20.3). Reference datasets were downloaded from the Stanford Large Network Dataset Collection (http://snap.stanford.edu/data/) and the BioPlex Interactome homepage (https://bioplex.hms.harvard.edu/interactions.php). The accompanying notebook is available as Supplementary File “Yeast_Network_comparisons.ipynb”. Gene annotation enrichment was performed using the 1D tool in Perseus (v.1.6.7.0). Annotation terms were filtered for 5% FDR (Benjamini–Hochberg correction) and a score above 0.

