## Supplementary Figures and Tables for "The social architecture of an in-depth cellular protein interactome"

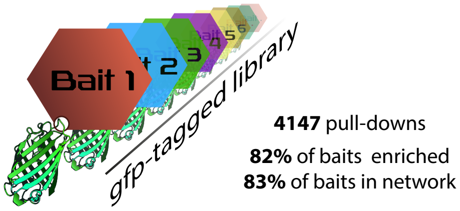


**Supplementary Figure 1.** Schematic of the GFP-tagged library. 4,147 different endogenous c-terminally tagged yeast strains (*7*) were used for 4,147 independent pull-down experiments. Each strain therefore allows the purification of the individually tagged protein (bait) and its specific interactors. The original library of 4159 strains was reduced by twelve strains to 4147, due to updates in ORF annotations (see methods: Data processing and normalization).

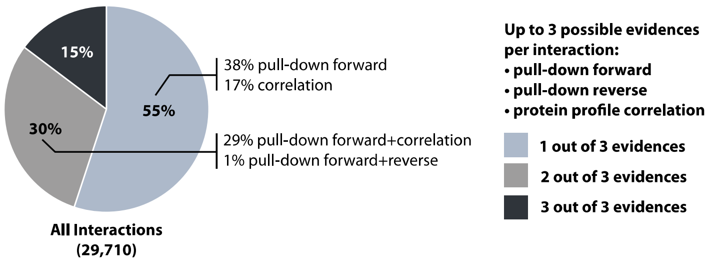


**Supplementary Figure 2.** Detailed proportion of interactions backed by multiple layers of evidence

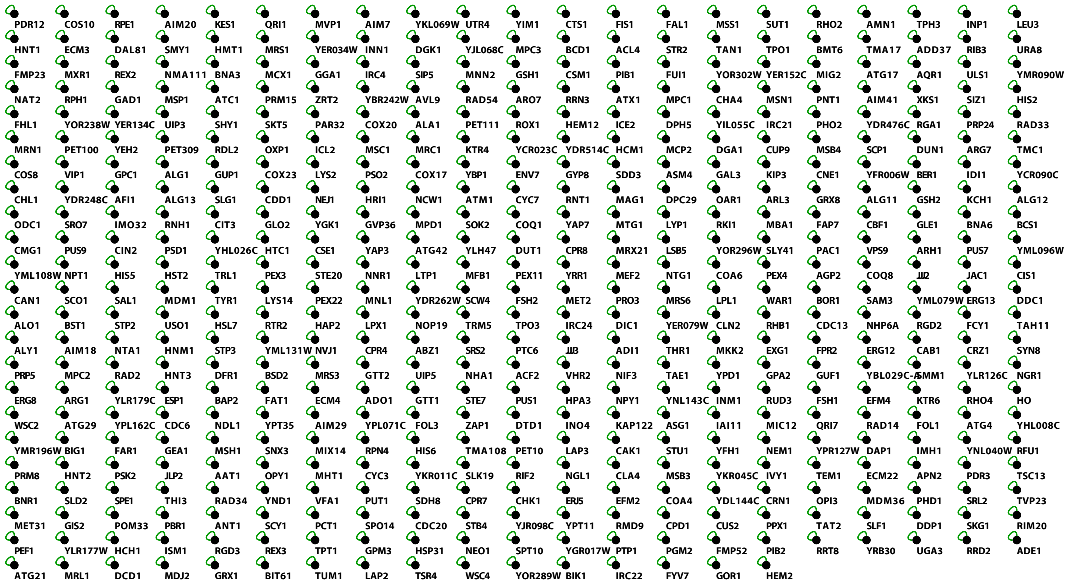


**Supplementary Figure 3.** “Asocial” proteins. Representation of 478 significantly enriched and detected bait proteins that lack any significant interactor under given conditions in this study. Green edges depict self-edges.

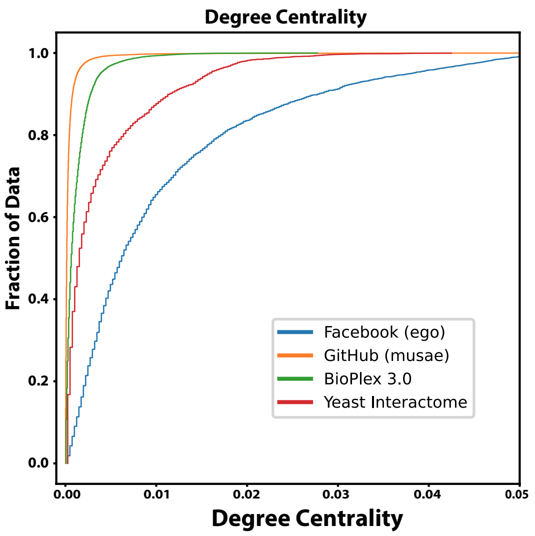


**Supplementary Figure 4**. Cumulative distribution function of the degree centrality. Comparison of different complex networks: *S. cerevisiae* has more influential (high degree centrality) nodes than BioPlex and GitHub, and less than Facebook.

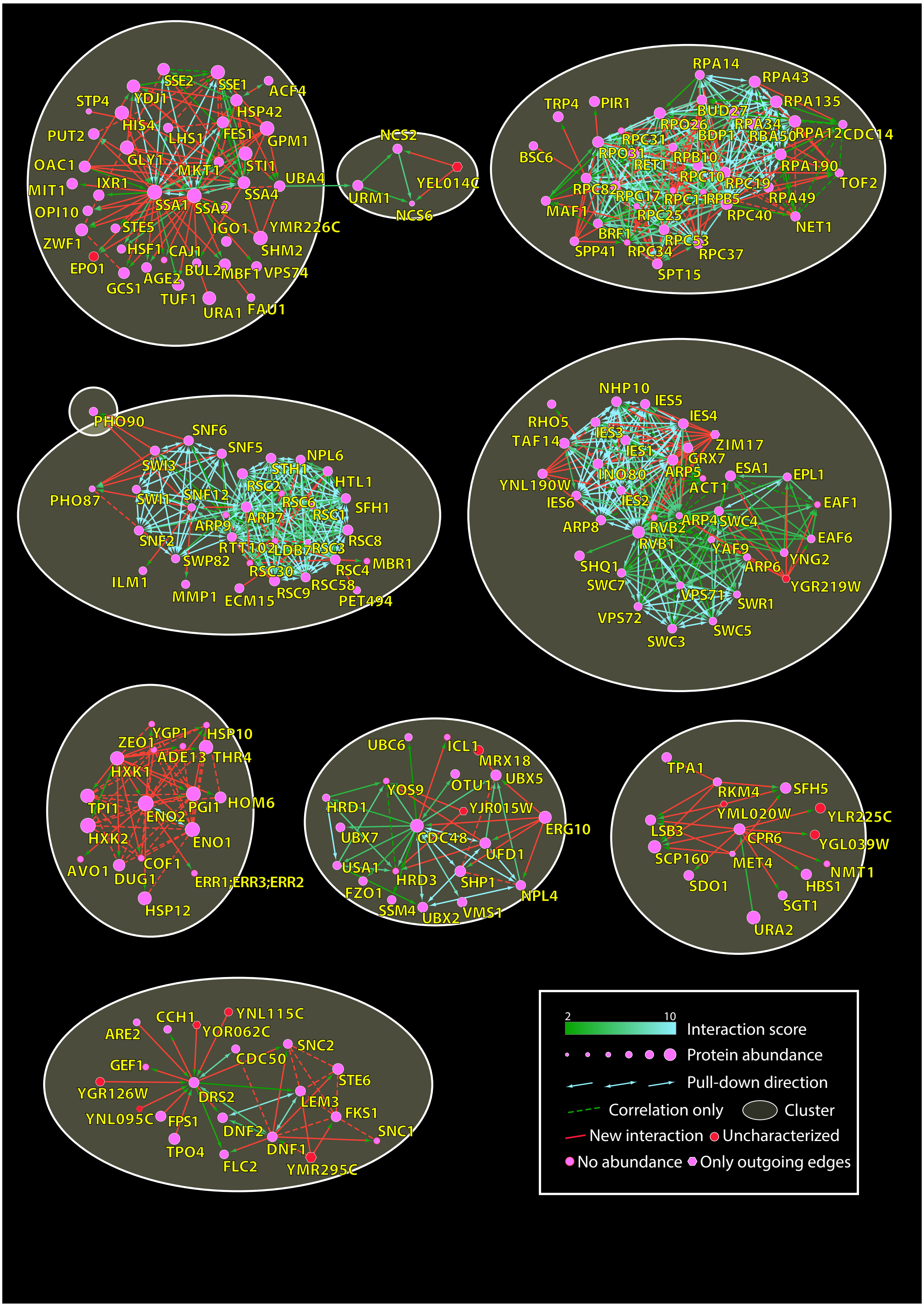


**Supplementary Figure 6 (part 1/5).** Extended selection of clusters involving proteins with novel interactions and/or uncharacterized proteins supported by multiple layers of evidence.

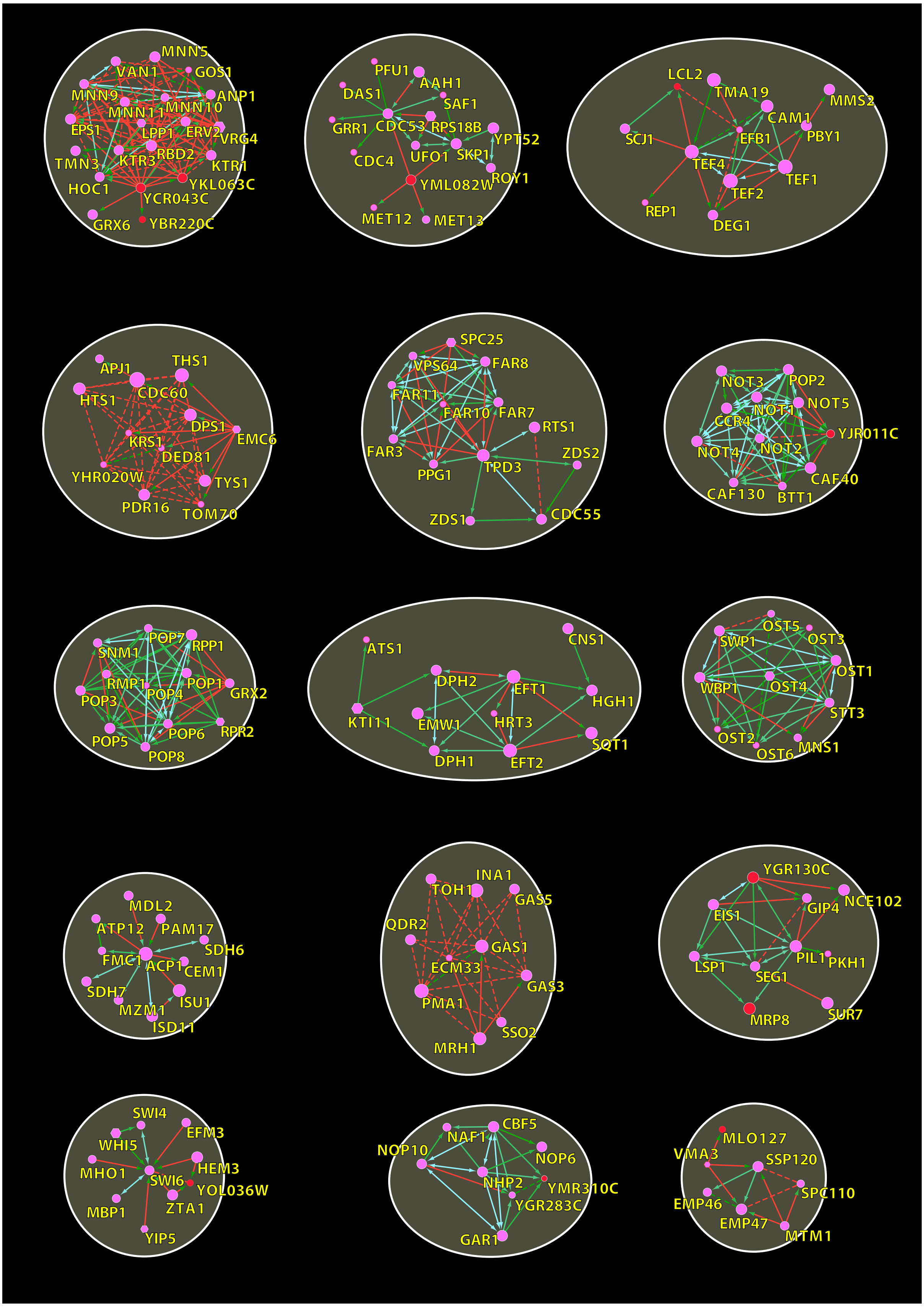


**Supplementary Figure 6 (part 2/5).** Continuation of Figure 6 part 2.

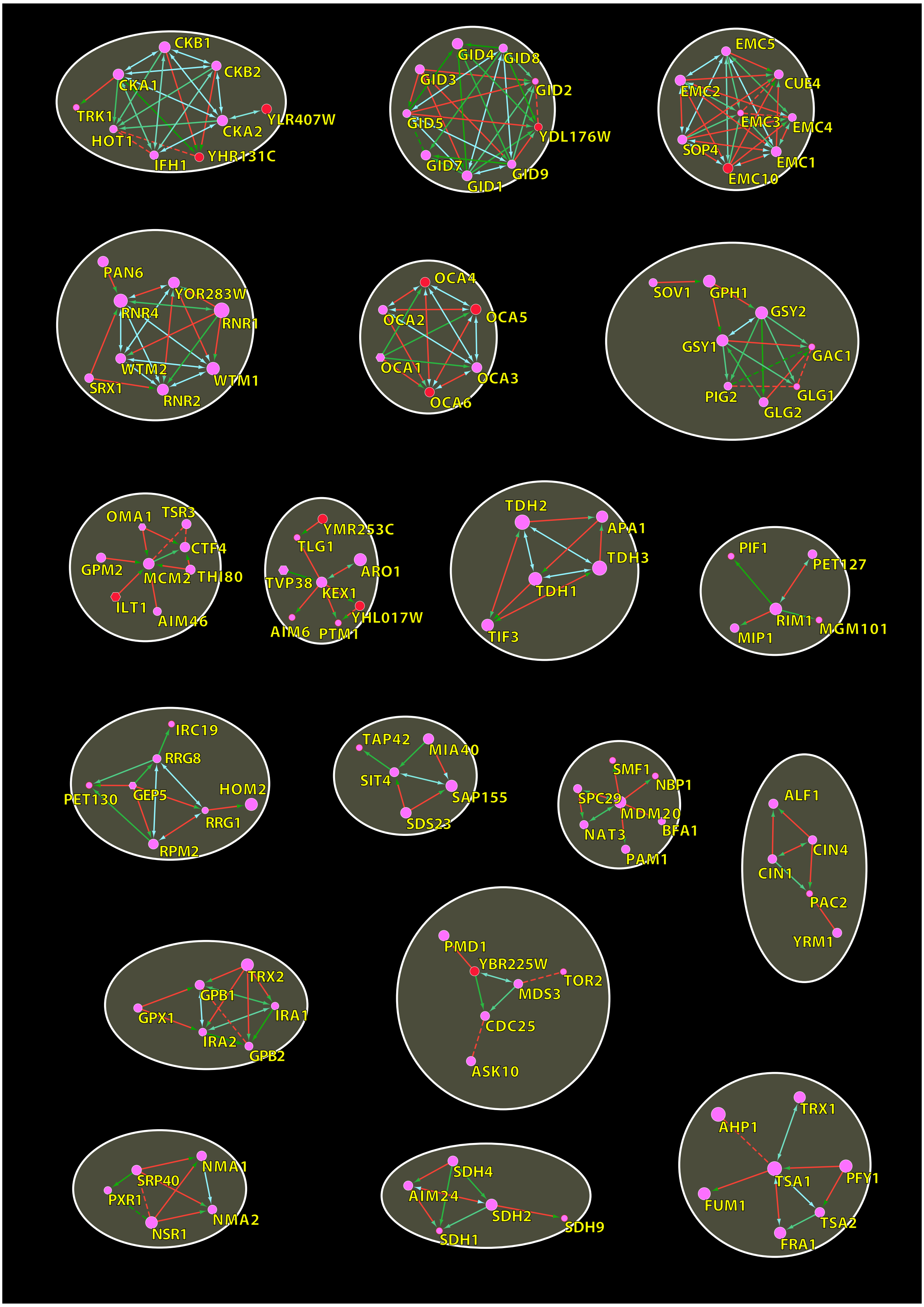


**Supplementary Figure 6 (part 3/5).** Continuation of Figure 6 part 3.

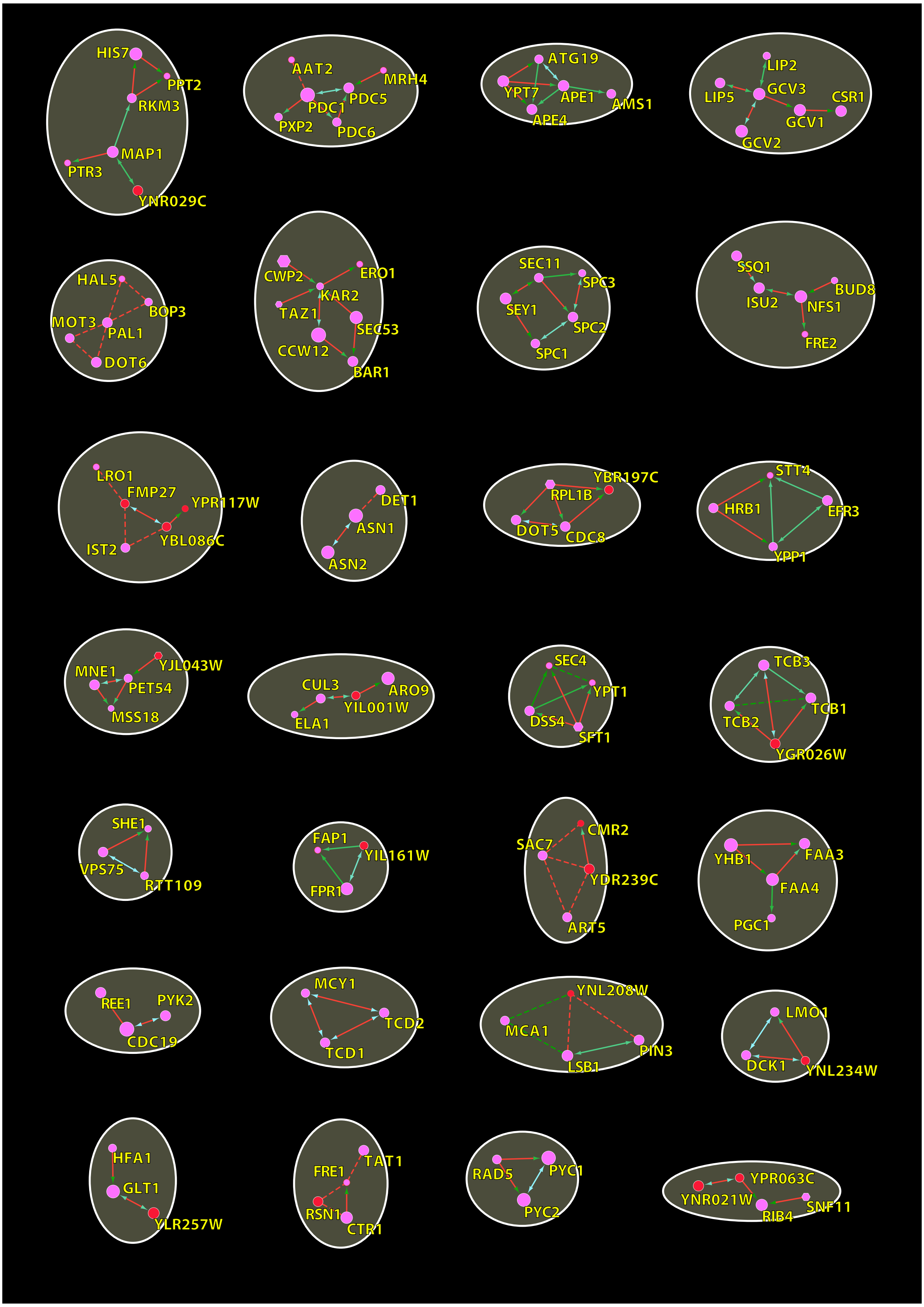


**Supplementary Figure 6 (part 4/5). Continuation** of Figure 6 part 4.

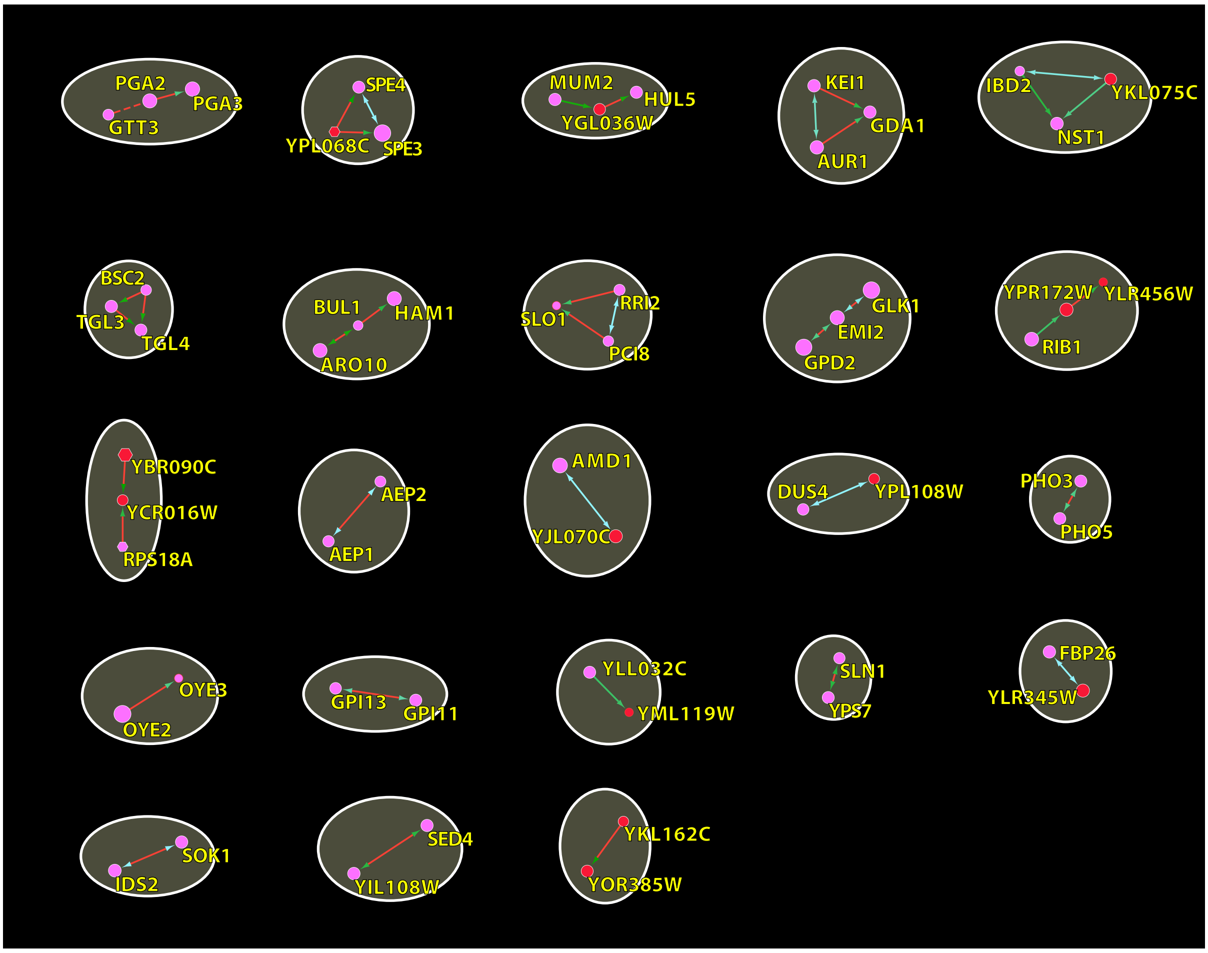

**Supplementary Figure 6 (part 5/5).** Continuation of Figure 6 part 5.

### Supplementary Tables

#### Table 1. Gene ontology term enrichment

| Gene ontology Name | Score | Benj. Hoch. FDR | -log10(p-value) | Size | Mean | Median |
|---|---|---|---|---|---|---|
| RNA polymerase II, core complex [GO:0005665] | 0,69 | 3,33E-02 | 1,64 | 11 | 2,49E-03 | 1,53E-03 |
| mitochondrial nucleoid [GO:0042645] | 0,67 | 3,71E-05 | 1,42 | 23 | 2,93E-03 | 1,30E-03 |
| gluconeogenesis [GO:0006094] | 0,64 | 3,77E-02 | 1,43 | 12 | 3,17E-03 | 1,47E-03 |
| misfolded protein binding [GO:0051787] | 0,58 | 2,99E-02 | 4,90 | 16 | 3,36E-03 | 1,33E-03 |
| polysome [GO:0005844] | 0,49 | 6,58E-03 | 1,44 | 28 | 2,48E-03 | 1,29E-03 |
| glycolytic process [GO:0006096] | 0,47 | 3,63E-02 | 1,46 | 22 | 3,42E-03 | 9,82E-04 |
| protein refolding [GO:0042026] | 0,45 | 4,25E-02 | 1,45 | 23 | 2,83E-03 | 7,32E-04 |
| proteasome storage granule [GO:0034515] | 0,44 | 3,89E-02 | 2,49 | 25 | 1,20E-03 | 8,86E-04 |
| ribosomal large subunit biogenesis [GO:0042273] | 0,39 | 3,47E-02 | 4,66 | 34 | 1,23E-03 | 8,15E-04 |
| cytoplasmic stress granule [GO:0010494] | 0,38 | 1,25E-05 | 1,48 | 82 | 2,11E-03 | 7,48E-04 |
| mitochondrial large ribosomal subunit [GO:0005762] | 0,35 | 2,31E-02 | 1,46 | 46 | 1,11E-03 | 6,90E-04 |
| preribosome, large subunit precursor [GO:0030687] | 0,30 | 2,20E-02 | 1,41 | 62 | 1,06E-03 | 3,53E-04 |
| mRNA binding [GO:0003729] | 0,26 | 2,17E-05 | 4,43 | 177 | 1,07E-03 | 5,23E-04 |
| ATPase activity [GO:0016887] | 0,25 | 2,16E-02 | 1,37 | 94 | 1,99E-03 | 4,71E-04 |
| mitochondrial translation [GO:0032543] | 0,23 | 3,45E-02 | 1,66 | 95 | 9,27E-04 | 4,33E-04 |
| RNA binding [GO:0003723] | 0,19 | 3,48E-04 | 1,52 | 273 | 1,15E-03 | 3,00E-04 |
| identical protein binding [GO:0042802] | 0,18 | 3,71E-02 | 3,46 | 156 | 9,40E-04 | 4,21E-04 |
| structural constituent of ribosome [GO:0003735] | 0,18 | 3,27E-03 | 2,18 | 242 | 8,29E-04 | 2,65E-04 |
| nucleolus [GO:0005730] | 0,16 | 3,57E-02 | 1,67 | 204 | 8,62E-04 | 2,74E-04 |

Gene ontology term enrichment on betweenness-centrality of nodes (proteins) in the network (1-dimensional annotation enrichment, FDR < 5%, score > 0).
